# Perpendicular Shear Stresses Drive Transmural Helical Remodeling in Engineered Human Ventricular Models

**DOI:** 10.1101/2022.08.18.504345

**Authors:** Nisa P. Williams, Kevin M. Beussman, John R. Foster, Marcus Rhodehamel, Charles A. Williams, Jonathan H. Tsui, Alec S.T. Smith, David L. Mack, Charles E. Murry, Nathan J. Sniadecki, Deok-Ho Kim

## Abstract

Tissue engineering with human induced pluripotent stem cell-derived cardiomyocytes enables unique opportunities for creating physiological models of the heart in vitro. However, there are few approaches available that can recapitulate the complex structure-function relationships that govern cardiac function at the macroscopic organ level. Here, we report a down-scaled, conical human 3D ventricular model with controllable cellular organization using multilayered, patterned cardiac sheets. Tissue engineered ventricles whose cardiomyocytes were pre-aligned parallel or perpendicular to the long axis outperformed those whose cardiomyocytes were angled or randomly oriented. Notably, the inner layers of perpendicular cardiac sheets realigned over 4 days into a parallel orientation, creating a helical transmural architecture, whereas minimal remodeling occurred in the parallel or angled sheets. Finite element analysis of engineered ventricles demonstrated that circumferential alignment leads to maximal perpendicular shear stress at the inner layer, whereas longitudinal orientation leads to maximal parallel stress. We hypothesize that cellular remodeling occurs to reduce perpendicular shear stresses in myocardium. This advanced platform provides evidence that physical forces such as shear stress drive self-organization of cardiac architecture.

## Introduction

With every contraction, the heart exhibits a unique pumping function where its muscle fibers shorten, thicken in diameter, and elicit a twisting motion of the whole organ^1^. The twisting is afforded by the change in orientation of myofibrils through the thickness of the myocardium^2^. Relative to the short axis or horizontal plane of the heart, the myofibers are orientated starting at -60° on the epicardial surface and shift to a +60° at the endocardial surface^3,4^. The twisting motion of the heart is like the wringing of a towel and is critical for efficient ejection of blood from the ventricles and therefore proper heart function^5^. When the mechanics of this motion are disrupted by disease or injury, heart function is compromised. For example, myocardial disarray is associated with several forms of cardiac disease (e.g. hypertrophic cardiomyopathy or infarction) and is thought to contribute to mechanical dysfunction^6–8^.

In addition to the heart’s distinct tissue architecture, its morphogenesis from an early structure that resembles a tube that folds and develops into a four-chambered organ has been studied for over two centuries^9,10^. Although animal models have provided a rich foundation of developmental biology from which we can glean great insight of human development, there is still a dearth of knowledge surrounding how the helical myocardial tissue patterning is developed. This knowledge gap is likely in part due to the difficulty of isolating and identifying biological governance over heart development in a complex whole-animal system. There is increasing evidence that mechanical cues may govern several aspects of heart morphogenesis^11,12^. However, there are few models with which these findings can be substantiated in the context of human biology^13^.

Tissue engineering strategies combined with human induced pluripotent stem cell (hiPSC) technology has enabled the development of diverse approaches for modeling structural and functional characteristics of cardiac tissue^14^. These efforts have provided complementary platforms to animal models for modeling human cardiomyopathies and drug cardiotoxicity testing in the dish^15^. However, most approaches yield two-dimensional (2D) laminar tissues or 3D structures that the lack structural complexity of the myocardium. There are few approaches that can recapitulate numerous aspects of the heart’s multi-scale organization within a single platform^16–20^, as it has been difficult to incorporate cardiomyocyte anisotropy and the 3D geometry of the ventricles into existing models. H. Chang *et. al* ^21^ has recently demonstrated the most advanced fabrication of multiscale tissue architecture by providing electrospun scaffolding with conical ventricular architecture and helical fiber orientations.

To address this gap of suitable technologies, we previously developed a cell-sheet engineering approach utilizing flexible thermoresponsive nanofabricated substrates (fTNFS) to enable production of 3D tissues with organized cellular architecture^22^. In this study, we adapted this platform to construct cardiac ventricular models with controlled architecture for studying the structure-function relationships within the myocardium. Given the functional significance of a heart’s highly organized, helical structure, we hypothesized that anisotropic tissue architectures would outperform isotropic ones due to their alignment of forces produced during contraction. To investigate this, we studied three structural organizations of the cell sheets within our 3D tissue models: circumferential (0°), angled (45°), and longitudinal (90°) cell orientations, and compared their contractile function to an isotropic control group with no cellular patterning. As hypothesized, the anisotropic ventricles outperformed the isotropic ventricles. Surprisingly, however, the circumferentially patterned models remodeled such that their inner layer realigned parallel to the chamber’s long axis. Finite element computational modeling revealed that this remodeling minimized transverse shear stress and maximized longitudinal shear stress. We postulate that cardiomyocytes are “sensitive” to shear stress along their short axis, which promotes cellular reorganization in favor of longitudinal shear. Our findings provide evidence for how mechanical forces might play a role in establishing the heart’s earliest helical tissue architecture.

## Results

### Fabrication of bioinspired tissue-engineered cardiac 3D ventricular models

To engineer miniaturized models of the human left ventricle, flexible thermoresponsive nanofabricated substrates (fTNFS) with nano-grooves and ridges were produced using soft capillary force lithography, as previously described^22^. The flexible TNFS were cut into fan shapes such that the nanogrooves were oriented at an angle of 90°, 45°, or 0° relative to the scaffold’s long axis to create longitudinal, angled, or circumferential organization, respectively (**Figure 1A & B**). Topographically flat scaffolds were utilized to create models with isotropic (random) cellular organization as a control. Each fTNFS was seeded twice with hiPSC-CMs and endothelial cells to form aligned cardiac sheets that exhibited coordinated spontaneous contraction patterns within five days of culture. The fTNFS was placed in a custom conical mold (**Supplemental Figure 1**), and a fibrin hydrogel (20 mg/mL) was cast in the interior to fabricate hollow ventricular models with final dimensions of 7 mm in height and 5 mm in diameter at the base (**Figure 1C-E**). Within one hour after removal from the mold, the isotropic, circumferential, and longitudinal tissues exhibited coordinated spontaneous contractions in which the apex was pulled upward and inward towards the base of the tissue (**Supplemental videos 1 & 2**). In contrast, the angled tissues exhibited an upwards twisting motion in the direction of the cellular patterning (**Supplemental video 3**). The coordinated spontaneous contractions of each tissue suggested the cell layers formed a functional tissue with coordinated contractions (**Figure 1E**). Histological analysis of the ventricular models indicated that the tissue’s wall thickness was approximately 320 µm, consisting of a 200-250 µm-thick fibrin wall encircled by layers of cells (~35-70 µm thick) (**Figure 1F and Supplemental Figure 2**).

**Figure 1.**
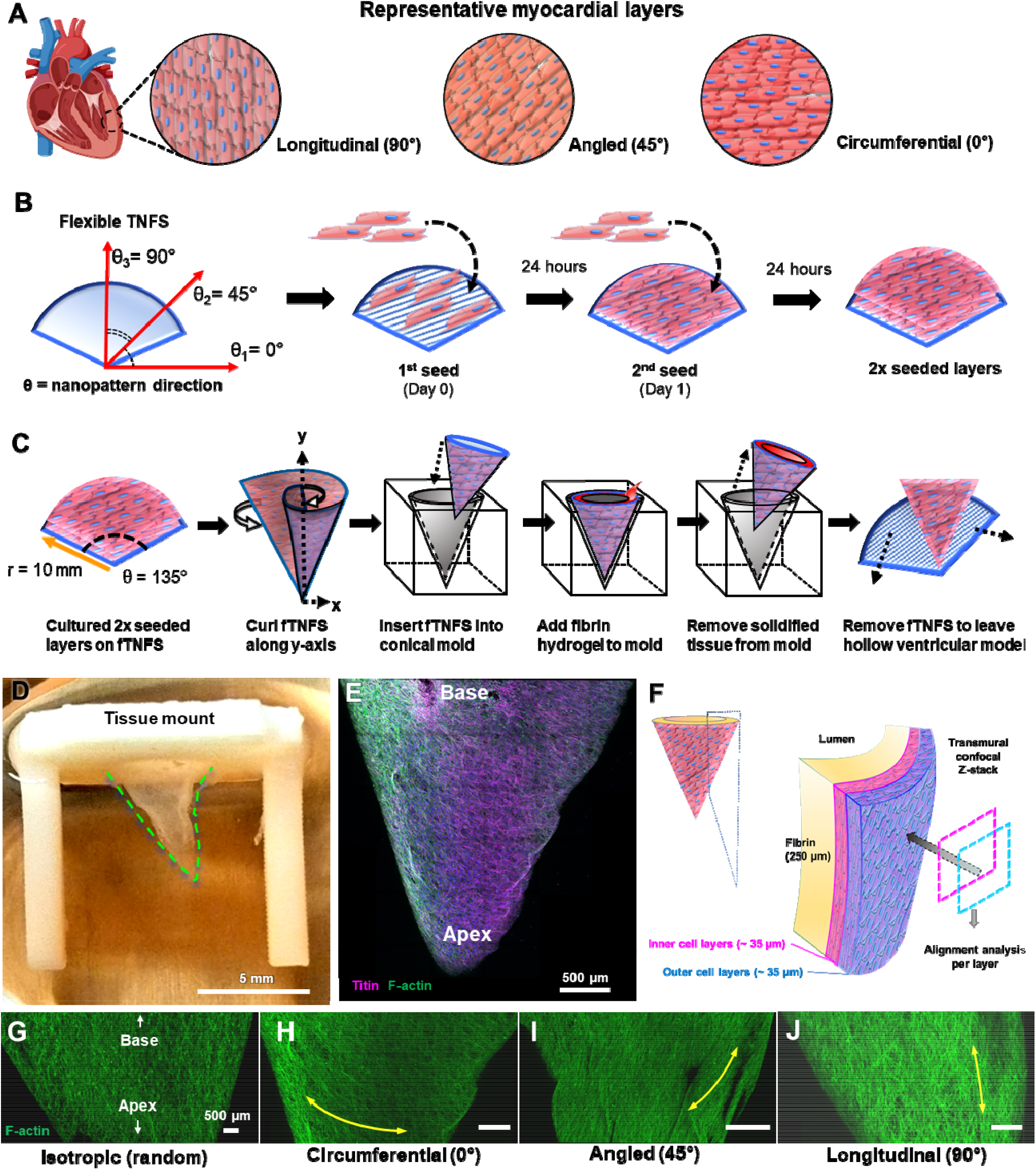
Design and fabrication of cardiac ventricular models. **(A)** Inspired by the layered organization of the myocardium, we chose to model three main cellular organizations in this study: longitudinal (90°), angled (45°), and circumferential (0°). **(B) (left)** Illustration of flexible **t**hermoresponsive **<u>n</u>**ano**f**abricated **<u>s</u>**ubstrates (TNFS) with direction of nanogrooves denoted (θ°). **(right)** Experimental timeline of serial cell seeding onto flexible TNFS for thick organized cell sheets. **(C)** Schematic of 3D ventricle model fabrication from organized cardiac sheets on fTNFS using custom molds and fibrin hydrogel **(see Supplemental Figure 1). (D)** Representative image of engineered ventricular model attached to a tissue mount in culture with tissue edge outlined in green dashed line. Tissues exhibited spontaneous contractions within one hour after removal from the molds **(Supplemental Video 1). (E)** 3D confocal z-stack projection of a circumferentially patterned ventricular model immediately after fabrication (day 0). The tissue was stained for sarcomeric protein titin (magenta) and filamentous actin fibers (F-actin, green). **(F)** Schematic of ventricular model cross-section highlighting inner fibrin wall covered by outer cell layers. **(right)** Overview of confocal imaging scheme to image through tissue wall from outer to inner cell layers. Each layer was analyzed separately for cellular alignment angle. **(G-J)** 3D confocal z-stacks of cardiac ventricular models fabricated with **(G)** isotropic (random), **(H)** circumferential (0°), **(I)** angled (45°), and **(J)** longitudinal (90°) cellular patterning. Cellular alignment is demonstrated by f-actin (green) organization in each model.

**Figure 2.**
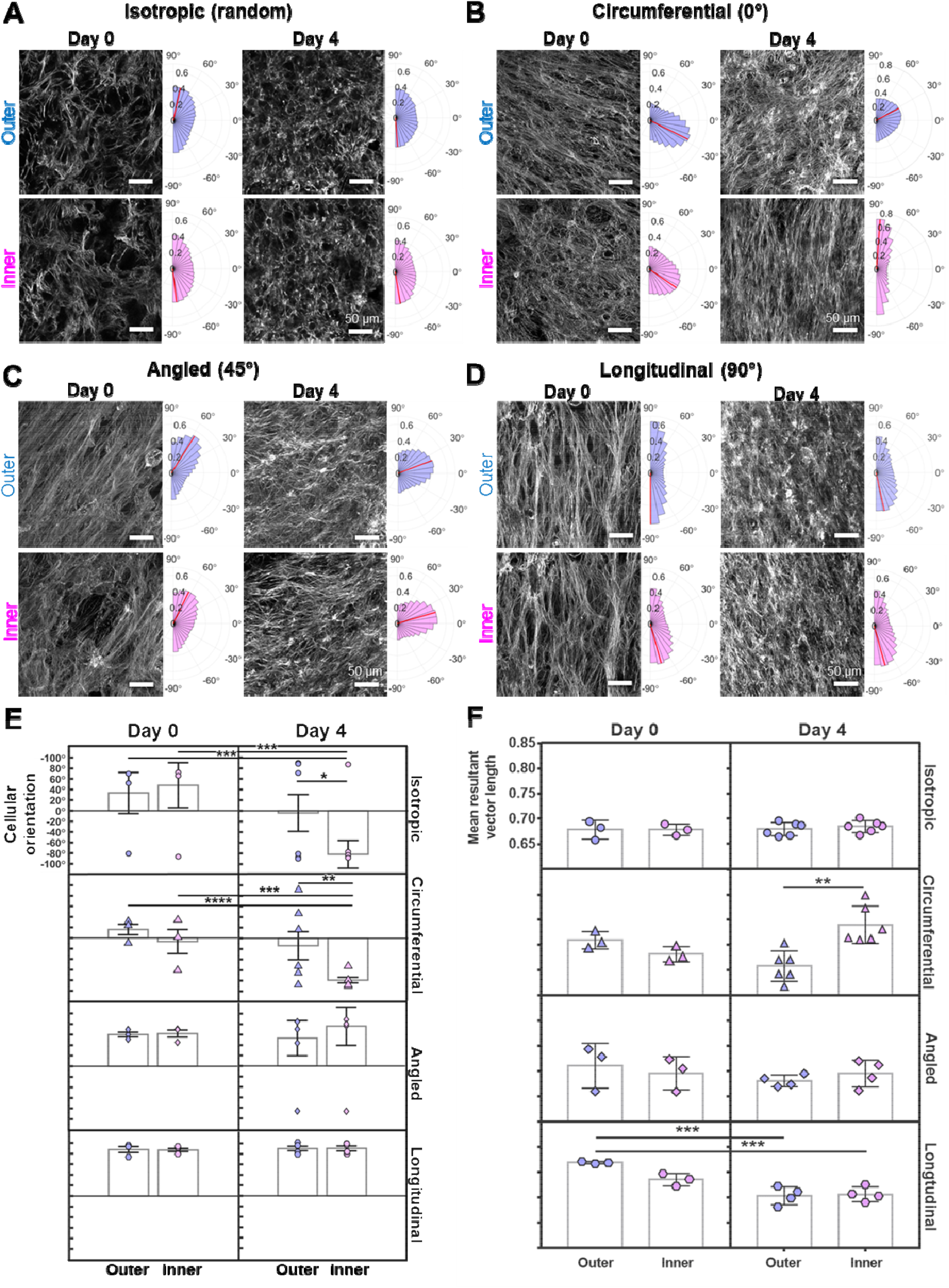
Quantification of cellular alignment in 3D cardiac ventricular models over time. 3D confocal z-stack projections of ventricular models patterned with **(A)** isotropic, **(B)** circumferential (0°), **(C)** angled (45°), and **(D)** longitudinal (90°) cellular organization as demonstrated by the filamentous actin organization (F-actin, grey). Representative images are from different tissues fixed and imaged on day 0 (immediately after fabrication) and day 4 of culture. All scale bars are 50 µm. **(Right of each image)** Polar histograms representing the distribution of cellular alignment of the inner (pink) and outer (blue) cell layers of each tissue organization (A-D), respectively. The height of each bar is the number of observations for that orientation angle relative to the total number of observations per image and is expressed as a fraction on the x-axis. Each histogram has 20 bins with a bin width of 9°. Red lines on each histogram represent the mean orientation angle for the specific image on its left but does not represent the mean of the group. **(E)** Average cellular orientation angles for outer (blue) and inner (pink) layers of each tissue group on day 0 and 4, respectively. **(F)** Average mean resultant vector length (RVL) for outer (blue) and inner (pink) layers of each tissue group on day 0 and 4, respectively. A RVL value closer to 1 can be interpreted as greater cellular organization towards the mean orientation angle for that tissue. Each data point in **(E & F)** represents the mean orientation angle of one tissue. **(E & F)** Error bars represent the standard error from the mean. *p < 0.05, **\***\*p < 0.01 ***p < 0.001, ****p < 0.0001.

### Cellular organization within 3D ventricular models is controllable

Immediately after the tissue casting process (day 0), ventricular models were fixed and stained to evaluate their microscopic cellular organization. High-magnification, confocal z-stacks were taken across the entire tissue area and transmurally through the cell layers. To determine if cellular organization was uniform, the outer and inner layers (~35 µm each, 2 cell-layers thick) of the z-stacks were parsed, analyzed separately, and compared. On day 0, the average cellular orientation for all cell layers for the circumferential (intended angle = 0°), angled (intended angle = 45°), and longitudinal (intended angle = 90°) groups were of 4.8° ± 7.4, 60.9° ± 3.2, and 87.1° ± 2.4, respectively (**Figure 2**). The isotropic group demonstrated some direction bias towards 40.8° ± 15.6, but the distribution of the cellular orientations was wide and lacked a prominent peak as compared to the other groups (**Figure 2E**). Across all groups, there was little difference between the angle of the inner and outer layers on day 0 (**Figure 2E, Day 0 column**). We next compared the mean resultant vector lengths (RVL) which describes the “intensity” of cellular organization around the average alignment angle^23^. For example, an RVL closer to 1 indicates a larger proportion of cells aligned in the same direction and an RVL closer to 0 indicates a lack of alignment (**Figure 2F, Day 0 column**). When comparing the RVLs of the outer and inner cell layers within each group, we found no significant difference between them at day 0. Thus, the intended cellular orientation of the cell sheets was maintained after the fabrication process for the ventricular models and uniform organization was maintained across both the inner and outer cell layers.

### Cellular remodeling occurs at the innermost cell layers of circumferential and isotropic tissues over time

To determine if cellular patterning in the ventricular models was maintained over the culture period, we fixed tissues after 4 days in culture and analyzed their cellular alignment (**Figure 2, day 4 columns**). In the isotropic group, only the inner cell layer remodeled, shifting its major axis from 48.2°± 42.4° to -81.6°± 26.4° SEM (p=0.00018), although there was no significant change in the mean resultant vector length (RVL) (**Figure 2A & E**). This suggested there was no increase in coordinated cellular organization towards the principal angle at -81.6° (**Figure 2F**). In contrast, tissues with circumferential patterning exhibited a distinct degree of remodeling over the culture period. The inner cell layers shifted their principal axis by 71.5° to become strongly longitudinally aligned (from -6.6°± 22.8 to -78.7°± 5.3, p = 0.0006), with no significant reduction in RVL values from day 0 to day 4 (**Figure 2E&F**). The outer cell layers shifted their major axis only slightly (from 16.2°± 9.9 to -14.1°± 26.4, p = 0.221) (**Figure 2E & F, Supplemental videos 4 & 5**). Tissues patterned with longitudinal or angled cellular alignment exhibited little change in their principal angle of alignment during the culture period for both the inner and outer cell layers (**Figure 2E**). For the angled tissue group, the RVLs were also not significantly different over time (**Figure 2F**). The outer layer of longitudinal tissues lost alignment strength compared to day 0, without changing the principal direction of organization. Thus, there was minimal structural remodeling in ventricular constructs with longitudinal or angled fiber orientation. Taken together, these data indicate that the inner cell layers of both circumferential and isotropic ventricular constructs remodel over 4 days by shifting their alignment toward a longitudinal orientation, whereas no remodeling was detected in either angled or longitudinal constructs.

### Cellular orientation contributes to engineered ventricle contractility

The characteristic helical tissue architecture of the myocardium results in a twisting motion during systole. Given the similar twisting movements exhibited by engineered ventricular models with angled (45°) cellular patterning (**Supplemental video 3**), we hypothesized that angled tissue patterning would functionally outperform all other tissue organizations which did not exhibit twisting motions immediately after fabrication. Additionally, we hypothesized that tissues with isotropic (random) cellular patterning would be functionally inferior to those with anisotropic tissue organization (e.g. circumferential, angled, longitudinal). To explore the relationship between cellular patterning and contractile performance, we evaluated each tissue organization for its ability to generate isovolumic pressures (**Figure 3A, B**). Pressure-sensing catheters were threaded into the lumens of each ventricular model after four days in culture and pressure production was recorded during electrically paced contractions at 1 Hz. Circumferentially and longitudinally patterned tissues generated 4 to 5-fold greater pressure amplitudes than isotropic tissues with random cellular organization (**Figure 3C**). The angled tissues generated an average of 3-fold greater pressure than randomly oriented tissues, but this did not reach statistical significance. The circumferentially and longitudinally patterned tissues contracted and relaxed 5-to 8-fold faster than the isotropic and angled tissues (**Figure 3D**). Although there were no differences in spontaneous beating frequency amongst the groups, the circumferentially and longitudinally patterned tissues had higher maximal capture rates in a ramped pacing protocol than the randomly patterned tissues (in circumferential and in longitudinal vs. isotropic, p<0.001; **Figure 3E-F**). These findings supported our hypothesis that anisotropic tissue organization promotes greater contractile function and also demonstrated that circumferential and longitudinal tissue organizations outperformed angled ones in our engineered ventricular model.

**Figure 3.**
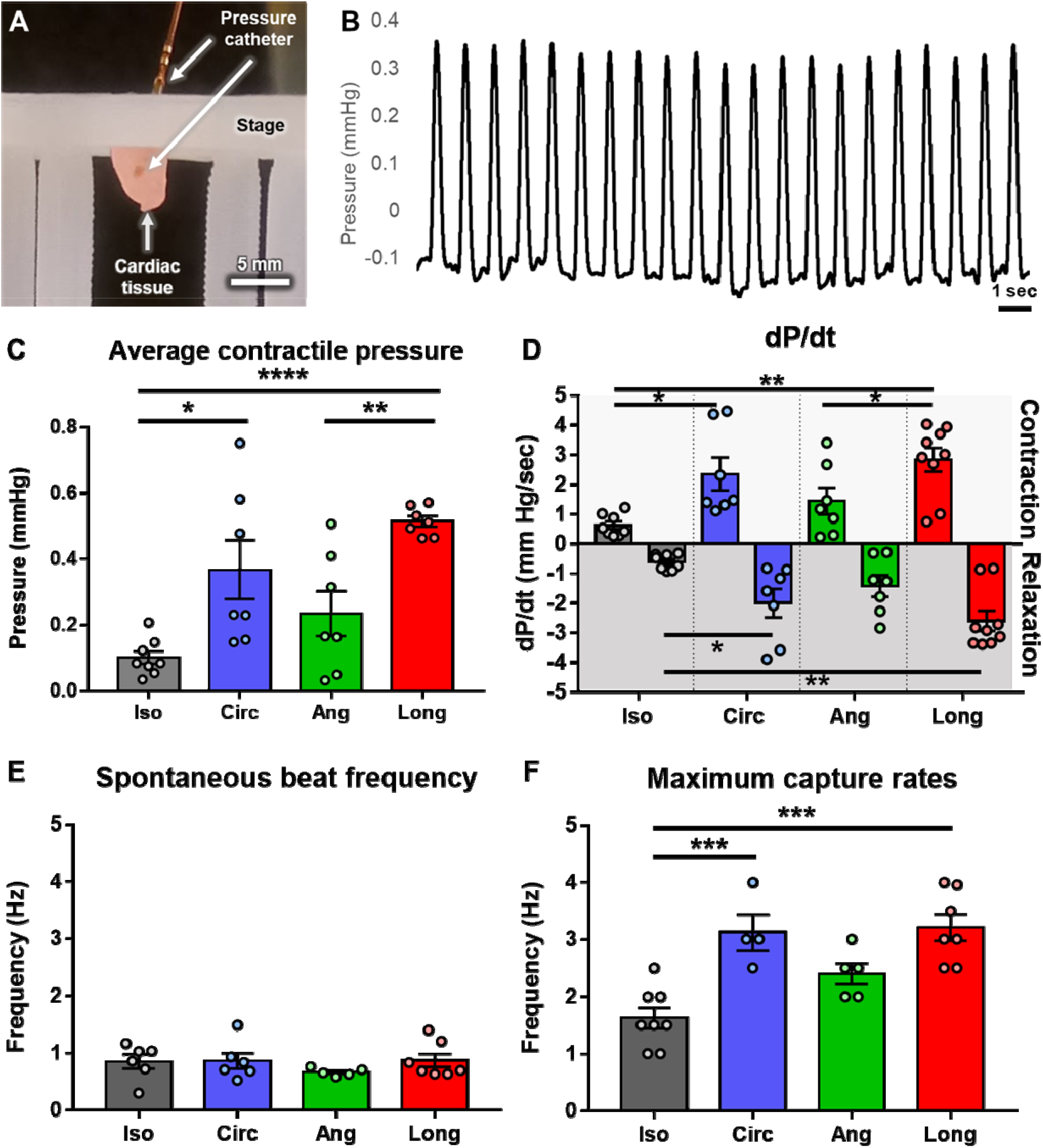
Functional assessment of ventricular models thorough isovolumic pressure production. **(A)** Image of ventricular model under catheterization during live pressure recordings. Tissues are positioned upright on a 3D printed stage during recordings. Culture medium was removed for a clearer view. **(B)** Representative pressure trace from a tissue under 1 Hz electrical field stimulation with 3 volts, 10 millisecond pulses. **(C)** Average contractile pressure amplitude from each tissue organization. Each data point represents one tissue within a group. **(D)** Average contraction (top) and relaxation (bottom) velocities for each tissue group as measured by the change in pressure over the change in time (*dP/dt*). **(E)** Average spontaneous (without electrical stimulation) beat frequencies recorded from several tissues within each group. **(F)** Average maximum pacing frequencies or capture rates for each group. **(C-F)** All measurements were taken on day 4 of tissue culture. Each data point represents average measurements from one tissue over a single recording. Error bars represent the standard error from the mean. * p < 0.05; ** p< 0.01; *** p < 0.001; *** p < 0.0001.

### Remodeling of the innermost layer reduces perpendicular shear stress

The striking remodeling effect that we observed in the circumferentially patterned tissues suggested to us that their cellular organization and therefore their contraction patterns may be providing mechanical cues that promote remodeling. We hypothesized that there might be unique differences in the transmural patterns of stresses and strains created by circumferentially patterned tissue contractions. Therefore, we next sought to model our system computationally to test our hypothesis and better understand the mechanical forces at play between tissues with different architectures. We built an axisymmetric finite element model to look at the stresses and strains that occur across the wall thickness in each of the conditional models at day 1 in culture. The model included the average tissue dimensions, stiffness of the fibrin hydrogel, wall thicknesses of the fibrin and cell layers, and the different cellular organizations (circumferential (0°), angled (45°), longitudinal (90°), and isotropic (random)) (**Figure 4A & B**). We first provided the model with experimental changes in tissue length from base to apex during contraction and relaxation observed in the longitudinally patterned tissues on day 1 (**Supplemental Figure 3, Videos 2 & 6**). This model then was used to predict shear stress and strain within the long and short tissue axes at any given point along the thickness of the tissue wall at peak systole for each pattern group (**Figure 4C & D, Supplemental Figure 4**). Shear stress and strain were studied and are reported here as they had relevance to the cellular realignment we observed in the case of circumferentially patterned constructs. We note that the short axis shear stress in the model was zero due to the annular structure of the model. Similarly, we considered hoop stress within the tissue wall, but this was zero through the fibrin wall and uniform throughout the cell layers. The apex of the model was free to move in space which also contributed to static hoop stress throughout cell layers. Since the experimental model was not a closed system during the culture period, pressure development within the chamber was not considered.

**Figure 4.**
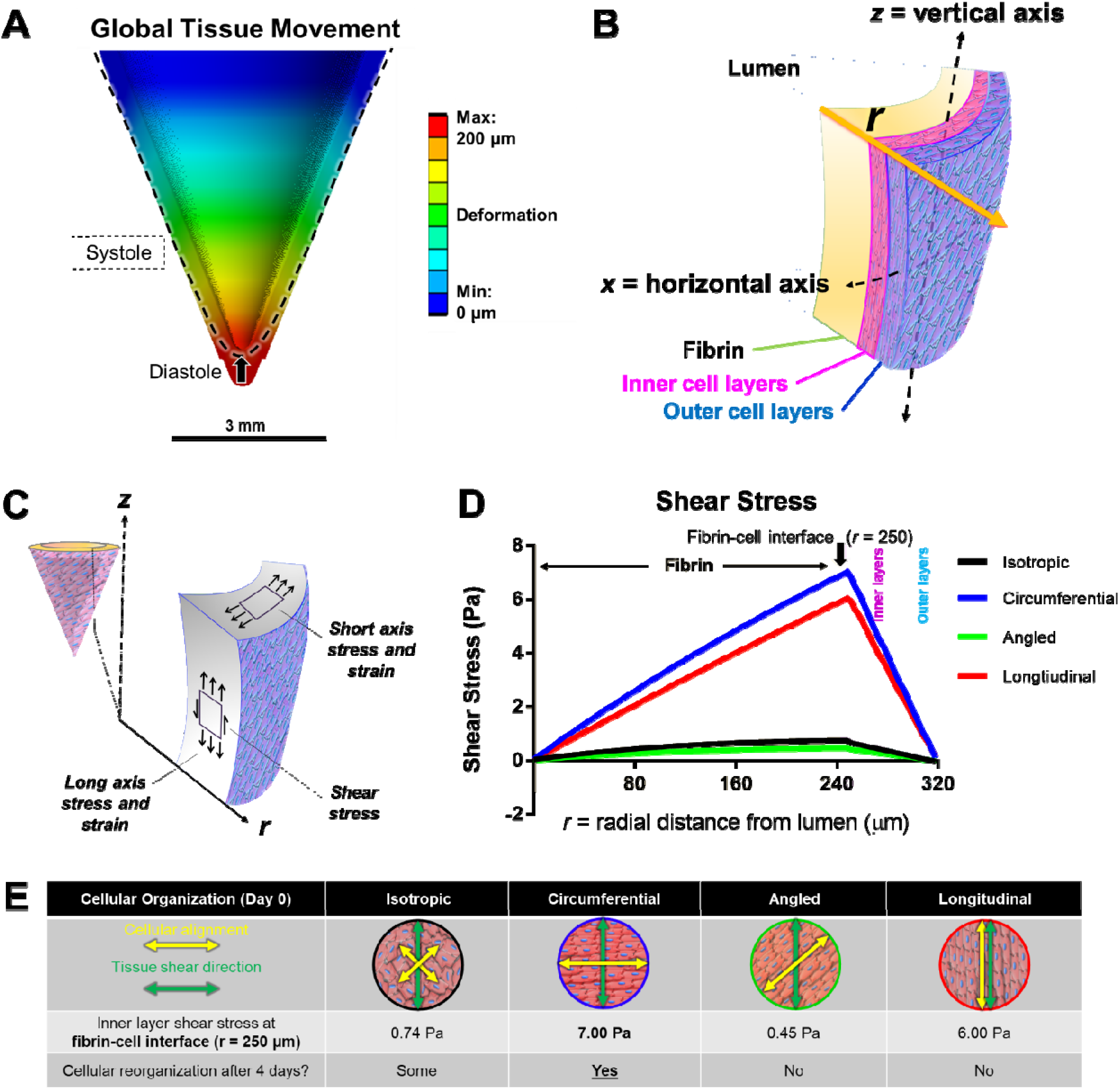
3D Finite element (FE) model of transmural shear stress. **(A)** 3D rendering of FE model of a longitudinally patterned tissue (cellular orientation = 90°). The color map projected onto the model represents the range of tissue deformation at peak systole. **(B)** Schematic of how transmural shear stresses were measured in **D and Supplemental Figure 4** where *r* = radial distance from the fibrin wall at the lumen space. Inner and outer cell layers are depicted in pink or blue, respectively. **(C)** Illustrations of the modeled shear stress and strains at any point along the tissue wall. **(D)** Shear stress measured in the vertical (*z*) axis direction across a tissue’s wall thickness (*r*) for all organizational groups. Yellow, pink and blue panels highlight the thickness and position of the fibrin wall, and the inner and outer cell layers, respectively. The fibrin-cell interface (at *r* = 250 µm) is marked with a black arrow. **(E, table)** Illustrations of the relationship between the directions of cellular alignment (row 1, yellow arrows) and vertical shear stress (green arrows) for each tissue group. The calculated vertical shear stress at the fibrin-cell interface (at *r* = 250 µm) is reported in row 2 for each tissue group. Indication of cellular remodeling that occurred during the 4 days in culture is reported in row 3.

From these simulations, tissues with isotropic or angled patterns were predicted to exhibit similar amplitudes and patterns of strain (i.e., tissue deformation) in both the long axis (apex to base) and short axis directions at any point throughout the tissue’s wall thickness (range: -2% to +2% strain) (**Supplemental Figure 3**). Similarly, isotropic and angled tissues had low levels of long axis shear stress across the thickness of the tissue wall at day 1 (range: -0.04 to +0.74 Pa) compared to the other experimental conditions which had a much wider range (range: ± 6.99 Pa) (**Figure 4D**). This suggested that isotropic and angled cellular organizations gave rise to almost equal tissue deformation in all directions and low shear stress in the long tissue axis where the major tissue movement was occurring from apex to base during systole (**Figure 4A**).

In contrast, tissues with longitudinal and circumferentially organized cells were predicted to produce the greatest amount of strain in the direction of their cellular alignment. Simply, circumferential tissues exhibit the greatest amount of short axis strain and longitudinal tissues exhibit the greatest amount of long axis strain (**Supplemental Figure 3**). For longitudinal and circumferential models, long axis shear stresses were maximal at the interface between the fibrin hydrogel and the innermost cell layers (**Figure 4D, *r* = 250 µm, black arrow**). In the case of circumferential models, the fibrin-cell interface was also where we observed the most pronounced degree of reorientation after 4 days in culture (**Figure 2B, E & F**). Although large shearing forces at the fibrin-cell interface were also predicted for longitudinally organized tissues, cellular remodeling was not observed in these tissues and cells remained aligned parallel with the direction of tissue movement in the long axis (**Figure 2D**). A critical distinction here is that for the circumferentially patterned tissues, the shear stress was perpendicular to cellular alignment, whereas for the longitudinally patterned tissues, the shear stress was parallel to cellular alignment (**Figure 4E**). This leads us to infer that cardiomyocytes realign to minimize shear stress that is perpendicular to their long axes. Perpendicular shear stresses were largest at the cell-fibrin interface and were associated with cellular remodeling towards the direction of shear along the long tissue axis. Furthermore, this example of cellular realignment towards the direction of high perpendicular shear stress was not observed in cell layers further away from the fibrin-cell interface (**Figure 4F**). These results suggest that cardiomyocytes are “sensitive” to shear stress along their short axis. Furthermore, the absence of remodeling in the outer layers suggests that there might be threshold of perpendicular shear stress needed for cellular realignment or that lesser shear stresses require longer times to induce remodeling.

## Discussion

The organization of the heart’s helical architecture results from concordant alignment of cardiomyocyte sarcomeric proteins, the cells within each myocardial layer, and the surrounding extracellular matrix, which together create the changing pitch angle between myocardial layers^1,5^. This multi-scale organization of the myocardium is critical for its proper pumping function, and there is still a minimal understanding of how this muscle pattern is generated during heart development^24^. There are few tissue-engineering approaches that can recapitulate multi-scale organization within a single platform. 3D bioprinting approaches allow for more precise placement of cells in space to create functional heart models with intricate chamber designs and directional fluid flow^17–19^. However, the cells in these models either are locally constrained by their printing matrix or lack organizational cues to encourage and maintain cellular organization on a global scale. Similarly, electrospun 3D fibrous scaffolds provide patterned extracellular matrix cues to align cells by providing single fiber orientations to guide cellular organization^16,25^. It is only very recently that electrospinning helical scaffold architecture has been demonstrated^21^.

Here, we addressed these limitations by incorporating nanoscale alignment cues to form highly organized cardiac sheets which were then casted into a 3D ventricular shape (**Figure 1**). This approach enabled us to generate cellular patterning on both a local and global tissue scale initially and then allowed cells to respond to mechanical cues generated by a more complex biomimetic environment. Our initial goal was to model different cellular organizations that exist within the myocardium and evaluate their structure-function relationships, testing the hypothesis that tissues with aligned cardiomyocytes would out-perform isotropic tissues. What we found was more complex. After four days in culture, circumferential and longitudinally patterned tissues mechanically outperformed isotropic and angled tissues and captured at higher pacing frequencies (**Figure 3**). As we explored the structural basis for this higher performance, we were surprised to learn that the circumferentially patterned tissues had remodeled their inner layers to a longitudinal orientation, whereas minimal remodeling occurred in the longitudinally patterned tissues (**Figure 2**). These results suggest that longitudinal tissue architecture, whether pre-patterned or spontaneously generated, was accompanied by greater pressure production, contraction and relaxation speeds, and faster pacing frequencies.

To gain a better understanding of forces that might drive selective remodeling of the circumferentially oriented tissues, we performed finite element analyses as the tissues moved upward and inward during systole (**Figure 4**). We found that circumferentially patterned tissues produced large, predicted shear stresses *perpendicular* to the cellular alignment on day 1 of culture. Conversely, longitudinally patterned tissues were predicted to produce large shear stresses *parallel* to the cellular alignment at day 1. We propose that perpendicular shear stress is a major driver to reorientation of the inner cardiomyocyte layer. In support of this idea, the magnitude of perpendicular shear was greatest at the fibrin-cell interface and decreased radially where the outermost cell layers were predicted to experience almost no shear (**Figure 4F**). This correlates with the transmural gradient of cellular reorientation, where there was a gradual shift in cellular alignment from 90° (longitudinal) at the inner cell wall, to near 15° (circumferential) at the middle layers, and almost random orientation at the outermost surface layer where the shear stress was predicted to be lowest (**Figure 2, Supplemental videos 4 & 5**). We interpreted these data to have three implications: (1) the initially circumferential contraction patterns produce great enough shear stress *perpendicular* to cellular alignment to promote cellular remodeling, (2) there is a “dose-dependency” of perpendicular shear stress required to promote remodeling by 4 days, and that (3) cells reorganize to minimize this high perpendicular shear by aligning parallel to its direction.

Cellular remodeling in response to mechanical stretching has been previously reported in other studies with adult rat cardiac fibroblasts^26,27^, endothelial cells^28–30^, smooth muscle cells^31^, dermal fibroblasts^32,33^; however, these studies were performed with either 2D cell monolayers or 3D laminar tissue patch settings. In the 2D settings, the cells often aligned their stress fibers perpendicularly to the principle direction of cyclic mechanical stretch as a form of “strain avoidance”^28–30^. However, when cells are embedded in a 3D hydrogel, the cells align parallel to the stretch direction ^34–37^. In our system, the cellular contraction forces provide their own mechanical stimulus and have a more complex 3D geometry than previous studies, so it is unclear which phenomenon is driving realignment. We speculate that strong shearing forces during contraction could “motivate” cellular organization away from or towards the direction of these forces.

These observations reminded us of the early embryonic heart tube which has a circumferential cellular patterning before it develops a helical pattern^38,39^. If we apply the beforementioned reasoning to the early heart tube, it seems plausible to think that similar transmural gradients of perpendicular shear stress could be evoked by the pumping motion of the tube against the cardiac jelly inside^40^. In this case, the cells closest to the cardiac jelly would begin to reorganize and align with the direction of shear. Meanwhile, the cells located in the outer layers would experience lower perpendicular shear and maintain their orientation, creating a pitch angle between the inner and outer layers. In effect, this gradient of perpendicular shear stress could explain how the initial helix muscle pattern is developed after the heart tube begins to beat.

Other mechanisms also may contribute to the heart’s twisting motion. For example, there is evidence that temporally spaced waves of contraction^24^ along the tube may contribute to the twisting motion required for its later asymmetrical C-looping when the heart begins to fold and form chambers. In our system, cardiac models were field stimulated such that all cells were contracting at once during systole, and we still observed remodeling. It would be interesting to apply apical point-stimulation to emulate the waves of contraction observed in the early heart tube and observe if greater remodeling from circumferential to helical is observed. Additionally, if supplementary circumferential cell layers could be added to an initial construct it might be possible that greater contraction force across more cell layers could inspire cellular remodeling into a more complete helix with a gradual shift in pitch angles between layers.

Overall, these results support the hypothesis that recapitulating the anisotropic organization of the myocardium within a physiological 3D tissue environment provide functional benefit compared to isotropic models. Additionally, the observed remodeling effect in response to gradients of transmural shearing forces has implications for how helical myocardial patterning might occur in cardiac morphogenesis and development. Our 3D ventricular model provides a powerful platform to recapitulate more complex myocardial tissue architecture and will allow for further investigation into human cardiac development.

## Materials and methods

### Fabrication of flexible thermoresponsive nanofabricated substrates (fTNFS)

To enable the production of cardiac ventricular models with anisotropic cellular organization, flexible films with nanoscale topographical cues and thermoresponsive properties were fabricated as previously described ^22,41^. Briefly, 100 µL of a UV-curable polyurethane acrylate (PUA) polymer (Norland Optical Adhesive #76) mixed with 1% w/w glycidyl methacrylate (GMA) was sandwiched and spread between a flexible polyethylene terephthalate (PET) film (5 × 5 cm) and a PUA master mold with nanoscale parallel ridges and grooves (800 × 800 × 600 nm, w x h x d). The PUA-GMA polymer mixture was flash cured under high-intensity UV light (365 nm) and the flexible film now with nanoscale features was removed from the master mold and placed under low-intensity UV bulbs overnight for final curing. The nanopatterned flexible films were then washed with an amine-terminated poly(N-isopropylacrylamide) (pNIPAM) solution (13 µM in H_2_O, M_n_ = 2500 Sigma-Aldrich) for 24 hours on a tabletop rocker (55 rpm, room temperature) to provide thermoresponsive surface functionalization. The flexible thermoresponsive nanofabricated substrates (fTNFS) were then rinsed in deionized water (DI-H_2_O) to remove excess pNIPAM).

Flexible TNFS were cut into fan shapes (radius = 12 mm, θ = 135°, area = 1.17 cm^2^) using a die cutter to produce substrates with nanogrooves aligned at either a 0°, 45°, or 90° angle relative to the x-axis (**Figure 1B**). Cut fTNFS were affixed to custom fan-shaped polydimethylsiloxane wells (PDMS, Sylgard 181) using a UV curable adhesive (Norland Optical Adhesive #83H). Wells were made only 10% larger in dimension than the fTNFS to control the cell seeding area and minimize superfluous cell waste. Culture wells were rinsed with DI-H_2_O before UV sterilization (294 nm) for 4+ hours in a biosafety cabinet. Sterilized culture wells were treated with fetal bovine serum (FBS, Sigma) overnight at 37°C before cell seeding to promote cellular attachment.

### Pluripotent stem-cell culture and differentiation

A human urine-derived induced pluripotent stem cell line (hiPSC UC 3-4, wild-type, male) was used for differentiation of cardiomyocytes (CMs) and endocardial-like endothelial cells (ECs) ^42^. Production of CMs and ECs was performed using well established monolayer-based directed differentiation protocols^43,44^. Briefly, hiPSC colonies were expanded to 80% confluency on Matrigel-coated plates (1:60, Corning), dissociated, and re-plated at either a high (270 k/cm^2^) or low (100 k/cm^2^) density for directed differentiation of CMs or ECs, respectively. High-density monolayers were cultured for 48 hours in mTeSR medium (STEMCELL Technologies) before induction (day 0) of mesoderm specification with 10 µm CHIR-99021 (Fischer Technologies) in Roswell Park Memorial Institute 1640 (RPMI) medium with B27 supplement without insulin (Gibco). To further specify cardiomyocyte lineage, high-density monolayers were exposed to the Wnt-inhibitor IWP4 (Stemgent) on day 3 in RPMI + B27 without insulin and cultured with RPMI + B27 with insulin from day 7 onwards. To purify differentiation populations for CMs, cardiac differentiated cultures re-plated and exposed to a glucose-poor and lactose-rich medium (RPMI 1640 without glucose or L-glutamine supplemented with 4mM lactate) on day 14 for two days or until only beating cells remained. Cells were harvested on day 17 or later and stained for cardiac-specific markers using a fluorescently conjugated antibody (anti cardiac troponin T (cTnT) – Alexa fluor 488, 1:100, Thermo-Scientific) for flow cytometry. Only populations of ≥ 95% cTnT-positivity were used for this study.

Endothelial cells were similarly differentiated from low-density (100 k/cm^2^) hiPSC monolayers plated in mTeSR medium with 1 µm CHIR-99021. After 24 hours, cells were induced with activin-A (R&D Systems) and Matrigel (1:60) in RPMI + B27 for 18 hours. The cells were then cultured with bone morphogenic protein-4 (BMP-4; R&D Systems) and CHIR-99021 in RPMI-B27 medium to specify for cardiac mesoderm lineages. To further select for cardiac endothelial populations, cells were incubated with StemPro-34 medium from days 2 to 5 with a cocktail of growth factors: vascular endothelial growth factor (VEGF; PeproTech), BMP-4, basic fibroblast growth factor (bFGF; R&D Systems), ascorbic acid, and monothioglycerol. On day 5, monolayers were dissociated and re-plated at a lower density (13k/cm^2^) on 0.1% gelatin-coated plates in Endothelial Growth Medium-2 (EGM-2, Lonza) supplemented with VEGF, bFGF, and CHIR-99021 until day 12. EC population purity was evaluated on day 12 via live-cell flow cytometry using an anti-CD31 – Alex 488 conjugated antibody (1:100, R&D Systems). Only EC populations with ≥ 90% CD31-positivity were used for this study.

### Serial-seeding of fTNFS with hiPSC-derived CMs and ECs

To eliminate confounding factors of cell age and maturation, CMs and ECs used between days 17-25 and 12-14, respectively. CMs and ECs were dissociated separately and mixed together such that final population was 89% CMs and 11% ECs (~7:1, CMs:ECs), as previously described^22^. The cell mixture was seeded onto FBS-treated fTNFS between 175 and 185 k/cm2 in 120 µL of cardiac growth medium (75% RPMI-B27 + insulin, 25% EGM-2, 10% FBS, 1% penicillin/streptomycin) to form a highly confluent monolayer (day 0). The cell mixture was cultured overnight at 37°C, 5% CO_2_ to allow for maximum cell adhesion to the fTNFS, mechanosensation of the nanotopography, and cellular elongation along the nanogrooves and ridges (**Figure 1B**). 18-24 hours after the first seeding event (day 1), additional CMs and ECs were dissociated and mixed again at a 7:1 ratio as described above. The excess medium and non-adherent cells were aspirated from the fTNFS surface and replaced with 120 µL of a second CM-EC cell suspension to provide another layer of cells between 175 and 185 k/cm^2^. The twice-seeded or serial-seeded fTNFS was cultured overnight at 37°C, 5% CO_2_ to allow for cell-cell adhesion to occur between the first and second seeded layers before addition of 2 mL of warmed (37°C) cardiac growth medium (day 2). Serial-seeded cell layers were cultured for an additional 4-5 days to allow for formation of aligned cardiac sheets with coordinated beating patterns before use in fabrication of 3D ventricular models.

### Custom molds for fabrication of 3D ventricle models

Modular 3D-printed molds were designed in a computer aided design software (Solidworks, Autodesk) and fabricated using a 3D-printer (CUBICON Style) and acrylonitrile butadiene styrene filament (Makerbot). The mold pieces were printed with a 0.1 mm line thickness and brushed with acetone before use to minimize the ridges formed by the layer-by-layer printing process. A modular design was incorporated to aid in mold disassembly and tissue extraction after fabrication (**Supplemental Figure 1**). The assembled molds contained a hollow conical lumen (Diameter = 6 mm, H = 7 mm) which was used to cast a tissue with a conical geometry that is like the left-ventricle of the heart. The final tissue product was on scale to a whole mouse heart^45^.

### Fabrication of 3D cardiac ventricular models from organized cardiac sheets

Organized cardiac sheets with spontaneous and synchronous beating patterns were formed after five days in culture on the fTNFS. Before tissue casting, 3D-printed mold pieces were pre-sterilized with 70% ethanol and submerged in hydrophobic Pluronic F-127 (5% in DI-water, Sigma) for at least 20 minutes to prevent the tissues from attaching to the molds. The submerged pieces were removed and allowed to dry in a sterile biosafety cabinet (BSC) for at least 5 minutes before assembly and tissue casting. Meanwhile, cardiac sheets were incubated with room-temperature phosphate buffer saline (PBS, Sigma) for 10 minutes to initiate partial cell sheet detachment from the fTNFS. The fTNFS was removed from the culture dishes using forceps to grasp both corners of the flexible pattern. Carefully, the two opposing corners were brought together and overlapped to create cone shape with the cell layers facing inwards (**Figure 1C**) and inserted into the complementary conical mold. One pair of forceps was used to hold the fTNFS in the mold while another was used to assemble the remaining pieces and fasten the fTNFS in place. To create a fibrin hydrogel scaffold, 18 µL of thrombin (50 units/mL, Sigma) was mix with 300 µL of fibrinogen (20 mg/mL, Sigma) and 200 µL of the thrombin/fibrinogen mixture was quickly pipetted into the open mold as to fill the entire well (**Supplementary Figure 1.iv**). Finally, the top mold piece was inserted through the mold’s opening and into the conical well, pushing excess fibrin out and causing it to flow into the remaining negative space of the mold. This overflow was essential for attaching the final casted tissue onto the tissue mount for future culture purposes.

The fully assembled mold containing the fTNFS and cell sheets was placed into a humidified 37°C incubator for 1 hour to allow for the thrombin/fibrinogen mixture to fully polymerize into a fibrin hydrogel scaffold within the mold. The top and bottom portions of the mold were then removed, and the remaining mold-fTNFS-cell sheet assembly was then submerged in cardiac growth medium and cultured overnight at 37°C, 5% CO_2_ to allow for the cell sheets to adhere to the newly polymerized fibrin hydrogel scaffold. After incubation, the mold was submerged in cold (4°C) medium and incubated at 4°C for 20 minutes to promote complete cell sheet detachment from the fTNFS. The mold was then fully disassembled and the fTNFS were removed leaving behind a hollow, ventricle-shaped tissue and organized cell sheets wrapped around the outside walls of the fibrin hydrogel scaffold (**Figure 1C and D**). The tissues were placed into 6 well plates with 9 mL of fresh medium for further culture.

### Tissue Culture and electrical field stimulation

After tissue casting (day 0) and extraction from the molds (day 1), ventricular models were cultured for an additional 24 hours before proving electrical field stimulation on days 2-4. On day 2, tissues were exposed to a 1 Hz pacing frequency (10 millisecond pulses, 3 V) for 24 hours and then increased to 1.5 Hz for an additional 1-2 days before functional measurements were taken on days 3-5 of culture.

### Functional assessment of ventricular models

After 4 days in culture, ventricular models were functionally evaluated using a pressure-sensing catheter. First, the ventricular models were transferred onto a custom 3D-printed stand within a 6 well-plate such that the tissues were positioned vertically with the base and the opening of the tissue lumen were at the highest point and the apex hung below. The wells were filled with warmed Tyrode’s solution (140 mM NaCl, 5 mM KCl, 5 mM HEPES, 1 mM NaH_2_PO_4_, pH 7.4) and a thin PDMS cover was placed over the opening of the tissue’s lumen at the base to create a closed-volume system. A small x-shaped slit was previously cut into the PDMS gasket to allow for the tip of a Millar pressure sensing catheter (model SPR-671) to be threaded into the lumen of the tissue (**Figure 5A**). Spontaneous pressure recordings were taken of each tissue to evaluate a baseline beat frequency using the Lab Chart Pro software (ADI Instruments). Platinum stimulating electrodes (□ = 0.5 mm) were then placed on either side of the tissue stand (15 mm apart) and the tissues were paced at 1 Hz with 10 ms pulses at 10 V. Pressure recordings were taken at 1 Hz for one minute before the pacing frequency was increased by 0.5 Hz and pressure production was recorded for another minute. This incremental pacing scheme was continued until the tissue could no longer capture at the challenging pacing frequency.

### Analysis of pressure production data

Spontaneous and electrically paced pressure recording events were parsed and exported from the LabChart Pro software as .csv files and imported into MATLAB (MathWorks) for analysis. A custom MATLAB script was used to find maximum peaks within each dataset and locate the preceding troughs to calculate the amplitude of each peak. The average pressure values at a 1 Hz pacing frequency for each tissue group (isotropic, patterned 0°, 45°, 90°) were averaged and compared using a one-way ANOVA and a Tukey’s multiple comparisons test (alpha = 0.05) (**Figure 5C**). Similarly, spontaneous and maximum capture rates for each group were averaged and compared using a one-way ANOVA and a Tukey’s multiple comparisons test (alpha = 0.05) (**Figure 5E & F**).

Tissues were also evaluated for their contractility through their ability to generate pressure over time during systole and diastole, or dP/dt. In the LabChart software, the first derivative of the raw pressure signal (dP/dt) was calculated for each tissue under each pacing frequency and averaged over the one-minute trace. The data were exported as a .csv file and imported into Prism where the average contraction and relaxation values for each group (isotropic, patterned 0°, 45°, 90°) were compared using a two-way ANOVA and Tukey’s multiple comparisons test (alpha = 0.05).

### Instron compression testing of fibrin

Compressive moduli of hydrated and crosslinked fibrin hydrogels (20 mg/mL) were measured using an Instron 5900 Series Universal Testing System equipped with a 10 N static load cell. Samples 5 mm in height and 6.8 mm in diameter were compressed at a rate of 10 mm/min until failure.

### Finite element analysis

An axisymmetric finite element model of the conical tissue was built in ANSYS to understand the realignment of tissues observed experimentally. The geometry shown in Figure 4 consisted of an inner fibrin layer 250 microns thick and an outer cardiomyocyte cell layer 70 microns thick. The inner diameter at the base was 5 mm and the length of the tissue from base to apex was 7 mm. A wedge-shape section of 2 degrees for the model was meshed with 32,026 nodes and 4,390 quadratic 3D solid elements (SOLID186 in ANSYS), and cyclic periodicity was applied with 180 repeats to model the full conical shape. Considering the small deformations of the conical tissues observed experimentally, an isotropic elasticity was applied to both the fibrin and cell layers with a Young’s modulus of 6.6 kPa and 20 kPa, respectively, and a nearly incompressible Poisson’s ratio of 0.49. Fibrin stiffness was determined experimentally through compression testing, as described above. To simulate contraction of the cardiomyocyte cell layer, a negative strain was applied to the cell layer causing either unidirectional or isotropic contraction. The direction of the contractile strain was rotated to replicate all experimental orientations, and the amount of contractile strain was calibrated by modeling the longitudinal orientation tissues and matching the apex displacement of the model to the one-day old experimental data (**Supplemental video 2**).

### Immunofluorescent staining and confocal imaging

Tissues were fixed in 4% paraformaldehyde for 24 hours at 4°C before immunocytochemistry was performed. Tissues were permeabilized in a phosphate buffered saline (PBS) solution with 0.2 % Triton-X 100 (Sigma-Aldrich, 9002-93-1), 5% goat serum, and 0.5% bovine serum albumen (BSA, Sigma-Aldrich A7906) for one hour at room temperature. After three, five-minute PBS washes, the tissues were incubated with an antigen blocking buffer (1.5% goat serum, 0.2% Triton-X 100) for two hours at room temperature to minimize non-specific antibody binding. Primary antibodies specific to sarcomeric titin (Myomedix, 1:300) were applied in a staining solution (0.2 % Triton-X, 1.5% goat serum) and incubated overnight at 4°C. Excess primary antibodies then were removed through three serial PBS washes before the secondary antibody (Alexa Fluor 647 donkey anti rabbit, 1:200) and fluorescently conjugated phalloidin (Invitrogen A12379, 1:200) were added. Tissues were incubated with secondary antibodies for two hours at room temperature in the dark before staining for nuclei DAPI (Invitrogen, D1306).

To aid in visualization of the large tissue surface area, the stained ventricular models were gently flattened and sandwiched between two glass coverslips using a thin (~3 mm) PDMS gasket with Vectashield anti-fade mounting medium (Vectro Labratories, H-1000-10). The tissues were then imaged using low- and high-powered objectives (20x air, 40x water immersion lens) and a SP8 Leica confocal microscope. Large-area stitched z-stacks were taken of the entire visible tissue area using the 20x-magnificaiton objective. For more detailed analysis of the cellular and cytoskeletal structure through the tissue walls, 40X-magnificaition z-stacks were taken at several locations across the tissue.

### Analysis of cellular organization

Several tissues within each group were fixed and stained at day 0 and day 4 of culture as described above. At least three confocal z-stacks were taken at 40x magnification across the tissue surface to survey transmural cellular alignment in the top (base), middle, and bottom (apex) sections. The z-stacks were parsed into halves (~35 µm thick) to separate out the outer and inner cell layers as defined by the nuclear counterstain (**Figure 1F**). These z-sections were collapsed to create maximum intensity projections (MIPs) representing the outer cell layers closest to the surface and inner cell layers closest to the fibrin wall of the tissue (**Figure 2**). Cellular alignment was determined for each of these MIPs based on alignment of the filamentous actin (F-actin) cytoskeleton using a modified MATLAB script as previously described^22,41,46^. Briefly, a low-pass Gaussian filter and edge detection to create a 2D convolution from which vertical and horizontal edges are detected using a Sobel filter^47^. These vectors are then used to calculate intensity gradient magnitudes across each pixel within an image. The images are processed by thresholding to define the edges of single cells or groups of cells and calculate their orientation angle between -90° and +90° relative to the x-axis at 0°. The total orientation angles detected within the image are binned and normalized using the probability density function in MATLAB (*normpdf*).

To quantify the principal alignment direction for the inner and outer cell layers of each tissue, the circular means for multiple images from each cell layer were calculated using the Circular Statistics MATLAB toolbox^23^. The mean orientations for the inner and outer layers of each tissue were then grouped and averaged (**Figure 2E**). The principal angles of alignment for each layer and tissue group were compared on day 0 (immediately after ventricle model fabrication) and day 4 using a parametric Watson-Williams multi-sample test for circular data (*circ_wwtest* function) and Tukey-Kramer multiple comparisons (alpha = 0.05).

To determine the strength of cellular alignment around the principal orientation, we calculated the resultant vector length (RVL) values for each image’s distribution using the Circular Statistics Toolbox (MATLAB, *circ_r* function). For example, if the distribution of angular data around the mean is very concentrated a RVL closer to one will result, whereas if the distribution is wide a RVL closer to zero will result. We interpreted RVLs closer to one to mean that the cells were highly aligned in the principal direction whereas RVLs closer to zero indicated the cells were less aligned in the principal direction. We compared average RVL values of the inner and outer layers within each group on days 0 and 4 using a one-way ANOVA with Tukey’s multiple comparisons (alpha = 0.05).

## Acknowledgements

We would like to thank the core facilities and staff of the Institute for Stem Cell and Regenerative Medicine (ISCRM), particularly the Tom and Sue Ellison Stem Cell Core and the Lynn and Mike Garvey Imaging Core. This work was supported by the National Institutes of Health: R01HL146436, UG3TR003271, R01HL156947, and R01HL135143 (to D.H.K.), NIH R01HL128362, R01 HL128368, R01 HL141570, and HL146868 (to C.E.M.) and 1F31HL145809-01A1 (to N.P.W.). The National Science Foundation grant CMMI-1661730 (to N.J.S.) also supported this work. Lastly, we are grateful for the institutional funding sources that supported this work: the Washington State funded ISCRM Fellows Program and the generous support from the Gree Real Estate company through the ISCRM Gree Scholars Program.

## Author contributions

<u>Nisa P. Williams</u>: conceptualization, methodology, validation, formal analysis, investigation, writing original draft, review and editing, visualization, funding acquisition

<u>Kevin Beussman</u>: methodology, software, formal analysis, writing original draft, review and editing

<u>Johnathan Foster</u>: investigation, review and editing <u>Marcus Rhodehamel</u>: investigation, review and editing

<u>Charles A. Williams</u>: data curation, software, review and editing

<u>Jonathan H. Tsui</u>: conceptualization, methodology, review and editing

<u>Alec S.T. Smith:</u> conceptualization, methodology, review and editing

<u>David L. Mack</u>: conceptualization, review and editing

<u>Charles E. Murry</u>: conceptualization, review and editing, supervision, resources, funding acquisition

<u>Nathan J. Sniadecki</u>: conceptualization, review and editing, supervision, project administration

<u>Deok-Ho Kim:</u> conceptualization, resources, review and editing, supervision, project administration, funding acquisition

## Supplemental Figures and Video captions

**Supplemental Figure 1.**
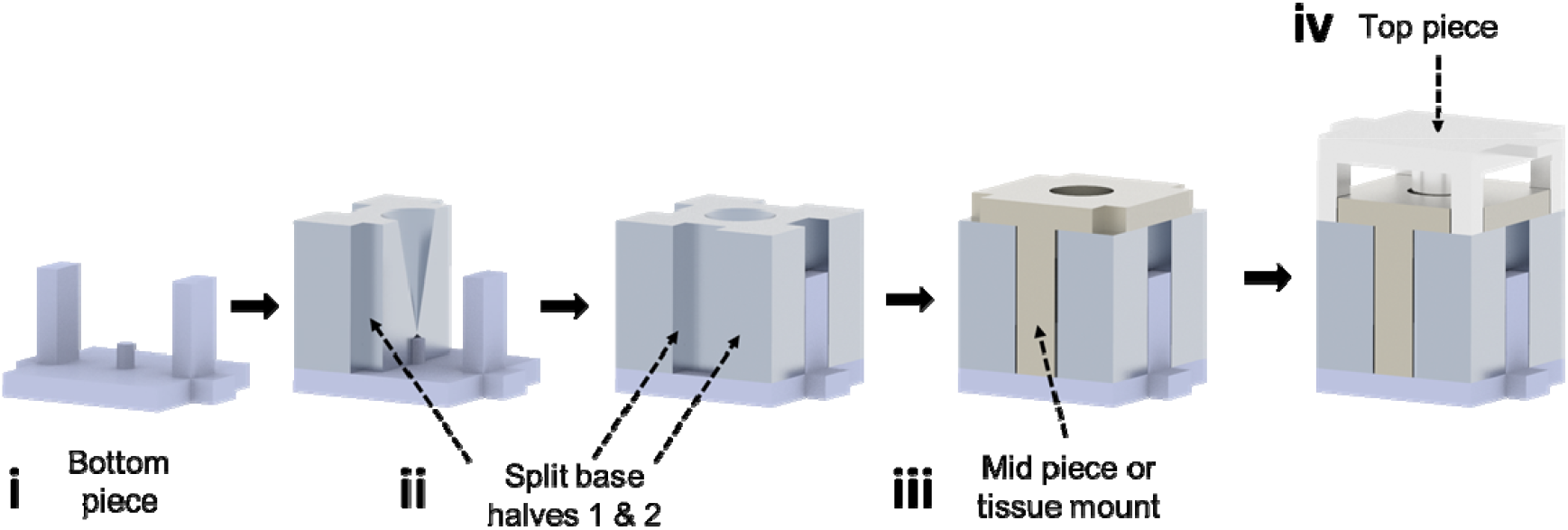
Assembly of custom conical molds for 3D ventricular tissue fabrication. 3D rendered models of tissue casting mold pieces and their assembly order. **(i)** The bottom mold piece is used as a platform to assemble the two base pieces **(ii)**. The base pieces join to create a conical lumen into which the fTNFS and cell sheets are folded and inserted into. A mid piece or tissue mount **(iii)** is inserted over top of the conical hole to prevent the fTNFS and cell sheets inside from springing out. The fibrin hydrogel is pipetted into the opening at this step. Finally, the top piece **(iv)** acts as a positive mold to create a hollow lumen in the final tissue.

**Supplemental Figure 2.**
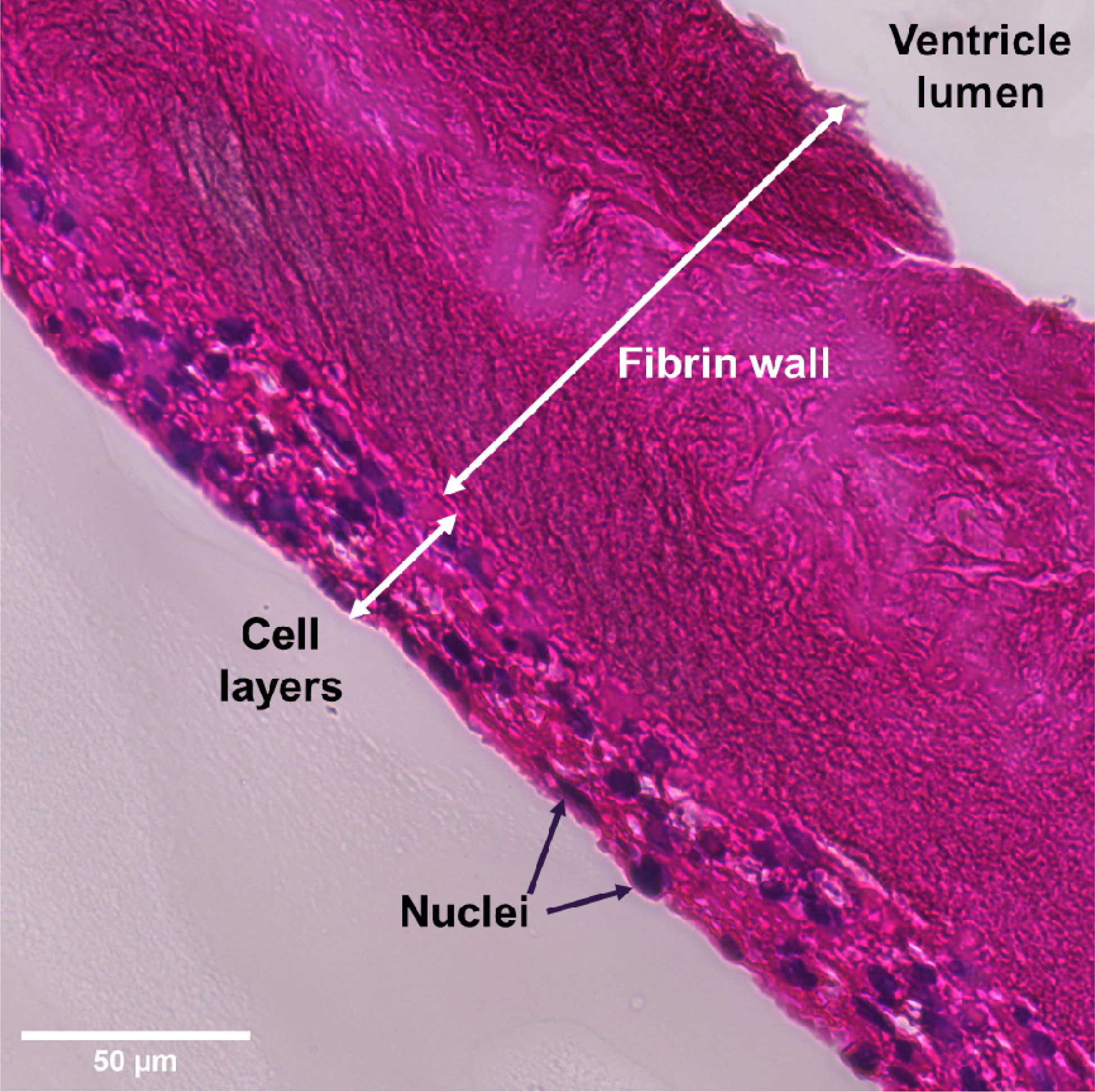
Long-axis cross section of an engineered ventricular model. Histological section of ventricular model stained with hematoxylin (nuclei, blue) and eosin (extracellular protein, magenta). Cell layers are attached to a thick fibrin wall that provides structural support and conical tissue architecture with a hollow lumen.

**Supplemental Figure 3.**
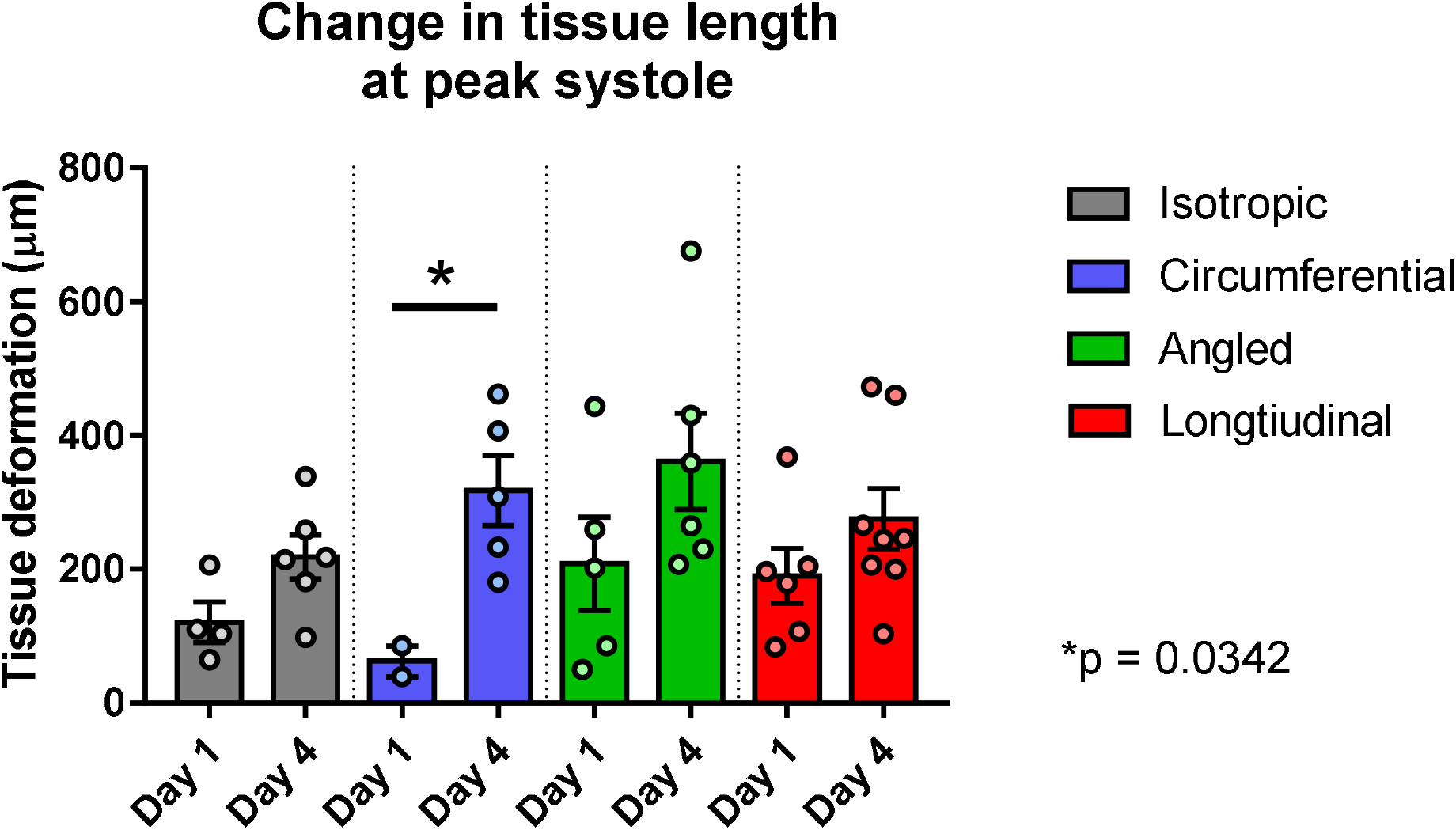
Total change in tissue length at peak systole. The length of each tissue from apex to base was measured at peak systole. Measurements were taken 1 day after tissue fabrication and at end point on day 4. The differences between tissues on days 1 and 4 were compared within the groups using a student’s t-test.

**Supplemental Figure 4.**
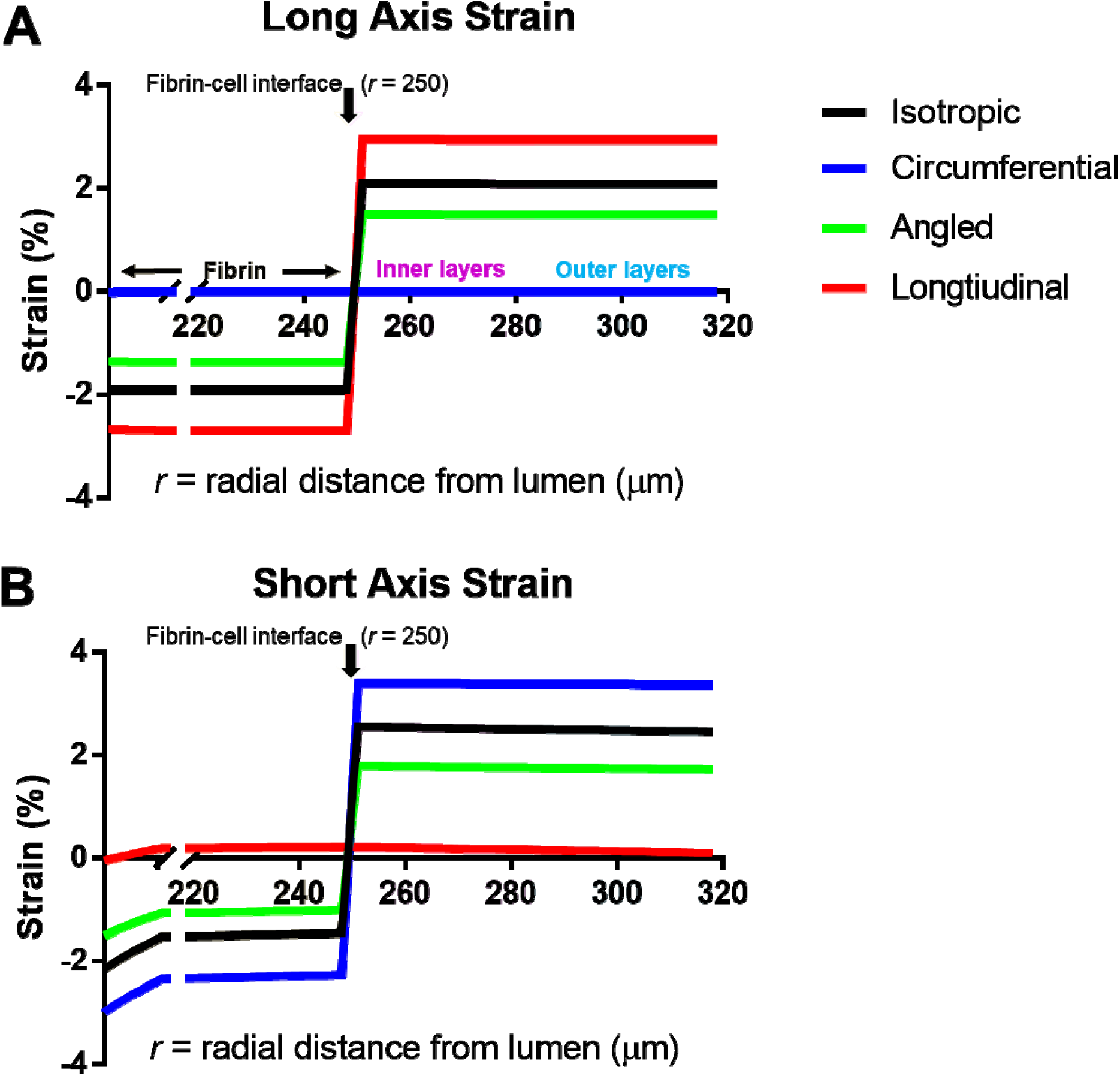
Finite element model analysis of transmural long and short axis strain.

**Supplemental video 1. Spontaneous tissue contraction after removal from casting molds**. Movie of spontaneous tissue contractions 1 hour after removal from the tissue casting molds detailed in main **Figure 1** and **Supplemental Figure 1**. The tissue is attached to a tissue mount during the casting process. The tissue mount is used to prevent the tissue from contacting the dish below and to allow for easy handling during functional measurements.

**Supplemental video 2. Longitudinal tissue movement during contraction and relaxation**. Movie of longitudinally patterned tissue (90° cell patterning) during stimulated contraction and relaxation after 1 day in culture with electrical field stimulation pulses paced at 1 Hz. The tissue movement during contraction produces an inward and upward movement towards the base of the ventricular model. During relaxation, the tissue extends back towards the apex. The overall tissue movement is about 387 µm. This inward and upward tissue movement was similar for the circumferential and isotropic tissues.

**Supplemental Video 3. Angled tissue movement produces twisting motion during systole. Angled tissue movement during contraction and relaxation**. Movie of a tissue patterned with angled (45°) cellular alignment one hour after removal from the tissue casting molds. Contraction along the angled cell alignment resulted in a twisting motion along the long axis of the tissue.

**Supplemental video 4. Circumferentially patterned tissues exhibit cellular remodeling over time**. 3D confocal z-stack movie of a representative circumferentially patterned tissue (day 0 organization = 0°) after 4 days in culture. The movie steps through the 70 µm of cell layers from outside to inside demonstrating the transmural change in angle of filamentous actin (F-actin, green) organization. Sarcomeric titin (red) and nuclei (DAPI, blue) are also labeled.

**Supplemental video 5. Sarcomere and thin filament organization changes transmurally in cultured circumferential tissues**. Confocal z-stack (70 µm total depth) through the wall of a circumferentially patterned tissue model after 4 days in culture. The movie steps through the 70 µm of cell layers from outside to inside demonstrating the transmural change in angle of filamentous actin (F-actin, green) and sarcomeric titin (red) organization.

**Supplemental video 6. Finite element model simulation of total tissue deformation of a ventricular model during systole and diastole**. The model simulates the movement of a circumferential or longitudinal ventricular model, as they have similar movement in our experimental models. The color scale depicts a minimum and maximum range of tissue deformation in microns.

